# Acute cold exposure in humans shifts the circulating proteome to a cardioprotective and anti-aging profile

**DOI:** 10.1101/2025.06.30.662436

**Authors:** Kaja Plucińska, Zsu-Zsu Chen, Ruijie Xiang, Samir Zaman, Lu Yan, Charlotte Hurson, Christian Peterson, Katie Fredrickson, Laurie Farrell, Jeanne Walker, Meltem Ece Kars, Gaurav Tiwari, Olivier Pourquie, Yuval Itan, Thomas S. Carroll, Kong Y. Chen, Aaron M. Cypess, Robert E. Gerszten, Paul Cohen

## Abstract

Cold exposure has been proposed to provide a constellation of salutary effects, yet its molecular correlates remain largely unknown. Brown adipose tissue (BAT) is the main site of adaptive thermogenesis, and its prevalence is linked with cardiometabolic health. Since the benefits of BAT activation and cold exposure more generally may be mediated through blood-borne factors, we conducted an extensive analysis of the circulating proteome linked with an acute cold challenge in healthy adults. Our goal was to uncover early molecular changes triggered by cooling and establish their specific relationships with the human brown adipocyte secretome as well as various phenotypic traits. Based on comprehensive inter-cohort validations, we provide the first reproducible proteomic signature of cold exposure in humans. Our data demonstrate that cooling favorably modulates circulating mediators linked with chronological aging, as well as metabolic and cardiovascular diseases, providing new potential biochemical transducers of the benefits associated with cold therapy.

**Highlights:**

- Cooling alters the plasma proteome with striking concordance in independent human cohorts.
- Cooling represses circulating proteins linked with type 2 diabetes, hypercholesterolemia, hypertension, coronary heart disease and heart failure.
- The circulating signature of cooling resembles a cardioprotective and anti-aging profile.

## INTRODUCTION

Cold exposure through modalities including ice water immersion or cryotherapy has garnered increasing attention for its proposed wide-ranging health benefits. While evidence supports the notion that cooling ameliorates inflammation, decreases fat mass and improves cardiovascular health^1, 2^, the underlying mechanisms are poorly understood. Enhanced sympathetic outflow to peripheral tissues during a cold challenge triggers a multi-organ response, including the activation of brown adipose tissue (BAT), a previously overlooked organ in adult humans^3–5^.

BAT functions as a “catabolic sink” for glucose, lipids and branched-chain amino acids, clearing these metabolites from the circulation, and performs uncoupled thermogenesis to convert chemical energy into heat in the cold^6, 7^. In mice, BAT activation is strongly associated with improved metabolic outcomes, including resistance to dietary obesity and improved glycemia, insulin sensitivity and lipid metabolism^8–10^. Substantially less is known about the benefits of BAT in humans and how they are mediated. Based on a propensity-matched cohort of over 50,000 patients, we reported that individuals with BAT have a 56% reduced odds of type 2 diabetes (T2D) and significantly lower odds of hypertension, dyslipidemia, and coronary artery disease, independent of body mass index (BMI)^11^. These findings were supported by lower blood glucose and triglycerides (TGs) in BAT positive (BAT+) individuals. The presence of BAT was further associated with decreased pathogenic visceral white fat and protection from hepatic steatosis^12^. Indeed, the beneficial effects of BAT overlap with metabolic biomarkers of extended lifespan^13^, but the responsible molecular mediators have been incompletely characterized.

Recent studies suggest that acute activation of BAT via mild cold exposure (14-19°C) or winter swimming is linked with elevated metabolic rate, higher energy expenditure (EE), lower fasting glucose^2^ and an improved plasma lipid profile in healthy men^14^. Repeated cooling interventions (4-6 weeks) further increase BAT volume, improve glucose and fatty acid (FA) uptake in skeletal muscle and reduce fat mass, even in individuals with low baseline BAT activity^15^, obesity^16^ or T2D^17^, though some of these benefits may be attributed to shivering-induced muscle thermogenesis^18, 19^. The metabolic benefits of BAT have generally been ascribed to its ability to convert chemical energy into heat. However, unlike murine BAT which accounts for about 1.5% of body weight, adult human BAT is estimated to constitute only 0.5% of total body mass and to consume at most 100 kcal daily^20–22^ in adult humans. These observations point to BAT-mediated pathways beyond energy expenditure that impact cardiometabolic health in humans.

Adipose tissue secretes an array of circulating factors termed adipokines, known to modulate blood glucose and insulin sensitivity (Adiponectin; RBP4; Resistin), β-cell function (Complement factor D, CFD) and food intake and energy expenditure (Leptin). Unsurprisingly, the secretory repertoire of BAT is distinct from white adipose tissue (WAT)^23–25^, given that the two tissues have different roles in energy metabolism. A growing body of literature has described BAT-derived secreted molecules, including lipid mediators, proteins, small peptides, and metabolites. Locally acting BAT-derived factors promote hypertrophy and hyperplasia of BAT and enhance innervation, and blood flow, aiding in BAT recruitment during increased thermogenic demand.

These so-called brown adipokines (or batokines) positively impact glucose metabolism (Slit2), liver fatty acid metabolism (NRG4) and immune responses (CXCL14). Embryonic BAT transplants also improved glycemia and WAT inflammation and modulated levels of adiponectin, IGF1 and FGF21 in mouse models of T1D and T2D^26, 27^, and substantially reduced visceral fat mass in genetically obese *ob/ob* mice^28^. Since the grafted BAT depots are small in size, it is conceivable that the disproportionate whole-body metabolic benefits are conveyed by soluble factors released from the donor tissue^27^.

In support of human batokines, 101 proteins were identified as uniquely secreted by primary human brown fat cells *in vitro*^23^. BAT activation also increases circulating levels of specific lipids and metabolites^29^, including 12-HEPE^30^ and 12,13-diHOME^9^. Translational studies have reported elevated plasma levels of both lipids in healthy human subjects post-cooling^14^, suggesting the cardiometabolic benefits of BAT may be mediated by bioactive, disease-modifying batokines. However, the field currently lacks a comprehensive atlas of circulating proteins regulated by cold exposure and BAT activation based on high throughput approaches and replication in independent cohorts.

Here, we provide a high-confidence, reproducible proteomic signature linked with acute cooling in healthy humans and identify associations between these factors and clinical traits related to body composition, BMI, cardiometabolic health, aging and major disease outcomes. We additionally studied the secretome of mature human induced pluripotent stem cell (hiPSC)-derived brown adipocytes and skeletal muscle cells *in vitro* to support cell type-specificity. This plasma proteomic signature can be leveraged to understand and harness the wide-ranging health benefits associated with cold exposure in humans.

## RESULTS

### Discovery and validation cohorts

To characterize the plasma proteome linked with acute cooling, we recruited 20 young, healthy subjects deemed highly likely to have cold-inducible BAT based on their age, BMI and lack of comorbidities (Discovery cohort; Fig. 1A). We collected blood at baseline (fasted > 12hrs; 22°C; BAT-inactive state) and after 3hr personalized cooling with a temperature-controlled water-perfused vest set 2°C above monitored shivering thresholds (ST) (fasted > 15hrs; BAT-active state). This cohort consisted of 10 males and 10 females of diverse ancestry, 21-28 years of age, with a BMI range of 18.5-25 kg/m^2^ (see anthropomorphic data in Table S1). Since subjects were fasted overnight, we reinvited this group for ‘fasting only’ blood draws to differentiate between effects of cooling versus fasting on the circulating proteome. We additionally studied an independent validation cohort of young, healthy men (n=12) and women (n=6) (Fig. 1A). This group had cold-induced BAT documented by ^18^FDG-PET/CT scans (Table S1), and was described previously^31, 32^. Subjects were on average 26 years of age, BMI 24.5, and for each, BAT status was characterized based on volume, activity and SUVMax^33^. Subjects had blood drawn at RT (22°C) and after 1hr of cooling (cooling vest set to 15°C). The female group had an additional 2hr cooling timepoint (Fig. 1A). To ensure reproducibility, we here focus on analytes that were significantly and concordantly changed with cooling in the two cohorts.

**Figure 1.**
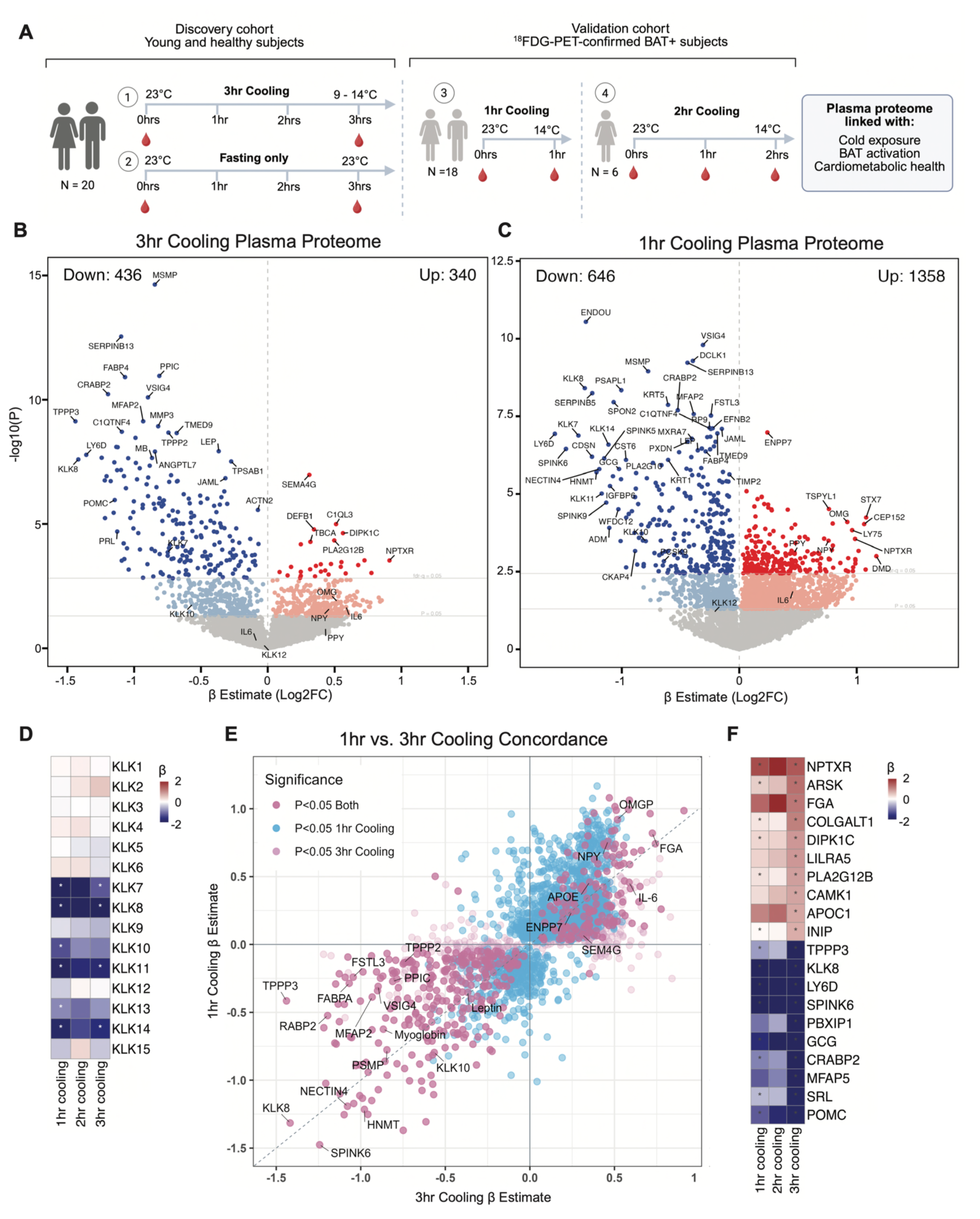
Acute cold exposure remodels the plasma proteome. **A.** Experimental design to identify changes in the circulating proteome in response to acute cooling in young and healthy individuals. For anthropomorphic data see Table S1. **B.** Effects of 3hr cooling on plasma proteome (*n* 19 individuals). See metadata in Table S2.1. For effects of fasting on the circulating proteome in the same subjects as well as plasma proteome validations see Figure S1. **C.** Effects of 1hr cooling on plasma proteome (*n* 16 individuals). See metadata in Table S2.5. **D.** Concordance of cold-regulated proteins across cohorts. Plotted are β-estimates for the 3hr (x axis) or 1hr cooling group (y axis). Proteins not regulated by cooling are not plotted. See metadata in Table S2.5. **E.** Effects of cold exposure on circulating Kallikrein (KLK) serine proteases (*n* 35 individuals). **F.** Top 20 cold-sensitive plasma proteins (FDR<0.05) concordant between cohorts (*n* 35 individuals).

### Acute cold exposure remodels the plasma proteome

To improve the analytical breadth of proteins we could measure, we used high-sensitivity, aptamer- and antibody-based platforms (SOMAscan and Olink; data from both platforms were merged, see Methods). We quantified 7,198 unique circulating proteins, of which 776 were significantly different between RT and 3hr cooling (p<0.05). 436 proteins decreased with 3hr cooling (p < 0.05; 225 by FDR < 0.05; Fig. 1B; Table S2.1), including hormones, paracrine signaling molecules, growth factors, enzymes and soluble receptors. These included adipose-derived proteins such as Leptin (β -0.37, p=1.19×10^-8^), CFD (β -1.07, p=2.58×10^-4^) and Fatty Acid Binding Protein 4, FABP4 (or FABPA, β -1.07, p=1.23×10^-11^), as well as Glucagon (β - 1.2, p=1.2×10^-7^), Lymphocyte antigen 6D (LY6D, β -1.36, p = 1.66×10^-8^) and Serine protease inhibitor Kazal-type 6 (SPINK6; β -1.24, p = 2.17×10^-8^). Leveraging control samples from individuals that returned for the non-cooling protocol, or ‘fasting only’, we established that 378 proteins were altered with fasting alone (p<0.05; Fig. S1A, Table S2.2), including Leptin (β -0.16, p=6.47×10^-2^) and FABP4 (cooling: β -1.07, p=1.23×10^-11^; fasting: β -0.16, p<0.05), both of which decreased with both perturbations. Although both cooling and fasting promote negative energy balance, many proteins were differentially regulated (Fig. S1B), including IL6 (cooling β 0.59, p=0.02; fasting β -0.98, p=0.01) and multiple Kallikrein (KLK) serine proteases. Notably, there were more proteins significantly suppressed with cooling in both cohorts (FDR<0.05, Fig. 1B-C). This leftward skew was associated with Cystatin C (CysC) levels, a proxy of estimated glomerular filtration rates (eGFR)^34, 35^, suggesting that cooling may have led to more rapid consumption, proteolysis, filtration, or dilution of plasma proteins. We therefore also included CysC-residualized data (Table S2.3 and S2.4, see also Fig. S1C). To independently validate the proteomic data, we measured several plasma proteins by ELISA and correlated protein concentrations (pg/mL-ug/mL) with SOMA and Olink data (Fig. S1D) and confirmed their statistically significant associations (p’s < 0.01).

### Plasma Kallikrein proteases are repressed with cooling independently of fasting

Interestingly, 16 serine proteases including Kallikreins, involved in proteolytic cleavage and activation of precursors linked with innate immunity, coagulation and cancer, were consistently downregulated after 3hr cooling and independent of fasting (Fig. S1B). These included KLK7 (3hr cooling: β -0.75, p = 1.38×10^-4^; fasting: β -0.32, p>0.05), KLK8 (3hr cooling: β -1.42, p = 2.53×10^-8^; fasting: β 0.02, p>0.05), KLK10 (3hr cooling β -0.57, p = 1.72 ×10^-2^; fasting β -0.52, p = 2.53×10^-8^, p>0.05), KLK11 (3hr cooling β -1.04, p = 6.82×10^-4^; fasting: β -0.02, p>0.05) and KLK14 (3hr cooling: β -1, p = 3.33×10^-6^; fasting β 0.39, p>0.05), while Kallikreins 1 to 6 were unchanged. Using the same discovery platforms, we detected 7,172 unique circulating proteins in the ^18^FDG-PET-confirmed BAT+ validation cohort (Fig. 1C). 2,004 of these proteins were significantly changed with 1hr cooling (p<0.05), and 518 passed the FDR cut-off (Table S2.5). Among the top cold-suppressed proteins were members of the Kallikrein family (Fig. 1D): KLK7 (1hr cooling: β - 1.37, p=1.32×10^-7^), KLK8 (1hr cooling: β -1.32, p=3.93×10^-9^), KLK11 (1hr cooling: β -1.17, p=9.7×10^-6^), and KLK14 (1hr cooling: β -1.11, p=2.58×10^-7^), as well as Leptin (1hr cooling: β -0.35, p=3.95×10^-7^), LY6D (1hr cooling: β -1.57, p=1.16×10^-7^) and SPINK6 (1hr cooling: β -1.48, p=3.55×10^-7^), similar to changes noted in the female cohort after 2hrs of cooling (Table S2.5).

### Cooling alters the circulating proteome with striking concordance in independent cohorts

We next directly assessed concordance between the two main cohorts (Fig.1E, Table S2.5). Despite differences in cooling duration, of the 225 cold-regulated proteins in the discovery cohort, 185 (81.5%) were also significant in the validation cohort (both FDR<0.05) with the same directionality. Twenty proteins increased and 165 decreased in the circulation after cooling, while the remaining 18.5% of proteins only reached significance in one of the cohorts or were differentially impacted by cooling duration. These data suggest that acute cold exposure promotes surprisingly similar changes in plasma proteomes across healthy subjects despite different cooling protocols. Of note, APOC1, NPTXR, FGA and ARSK were amongst the most upregulated with cooling, while KLK8, LY6D, Pro-opiomelanocortin (POMC), and SPINK6 were among the most downregulated factors (Fig.1F).

### Smooth muscle and adipose-derived proteins exhibit larger changes with cooling

Since acute cold exposure induces changes across many tissues, we next aimed to elucidate the potential source of cold-regulated plasma proteins by mapping all plasma data including the concordant 254 proteins onto the Human Protein Atlas (HPA)^36^, which contains bulk RNA-Seq data from 50 organs. Here, we considered a protein to be enriched in a specific tissue if its expression was 4-fold higher than its mean expression across other organs. We then took all of the proteins detected in the plasma in our study and determined how many of them showed tissue-specific enrichment. Interestingly, of 42 proteins that mapped to smooth muscle 23 of them were cold-responsive (55%), making this the tissue with the highest percentage of change. Next, 49% of adipose-mapped proteins changed with cooling (71 of 144), while only 35% of skeletal muscle and 33% of liver-derived proteins were altered with cold (Fig. 2A). Among the adipose-derived proteins were Leptin, CFD, FABP4, lipoprotein lipase (LPL), angiopoietin-like protein 4 (AGPTL4), V-Set Immunoglobulin Domain Containing 4 (VSIG4) and NNMT.

**Figure 2.**
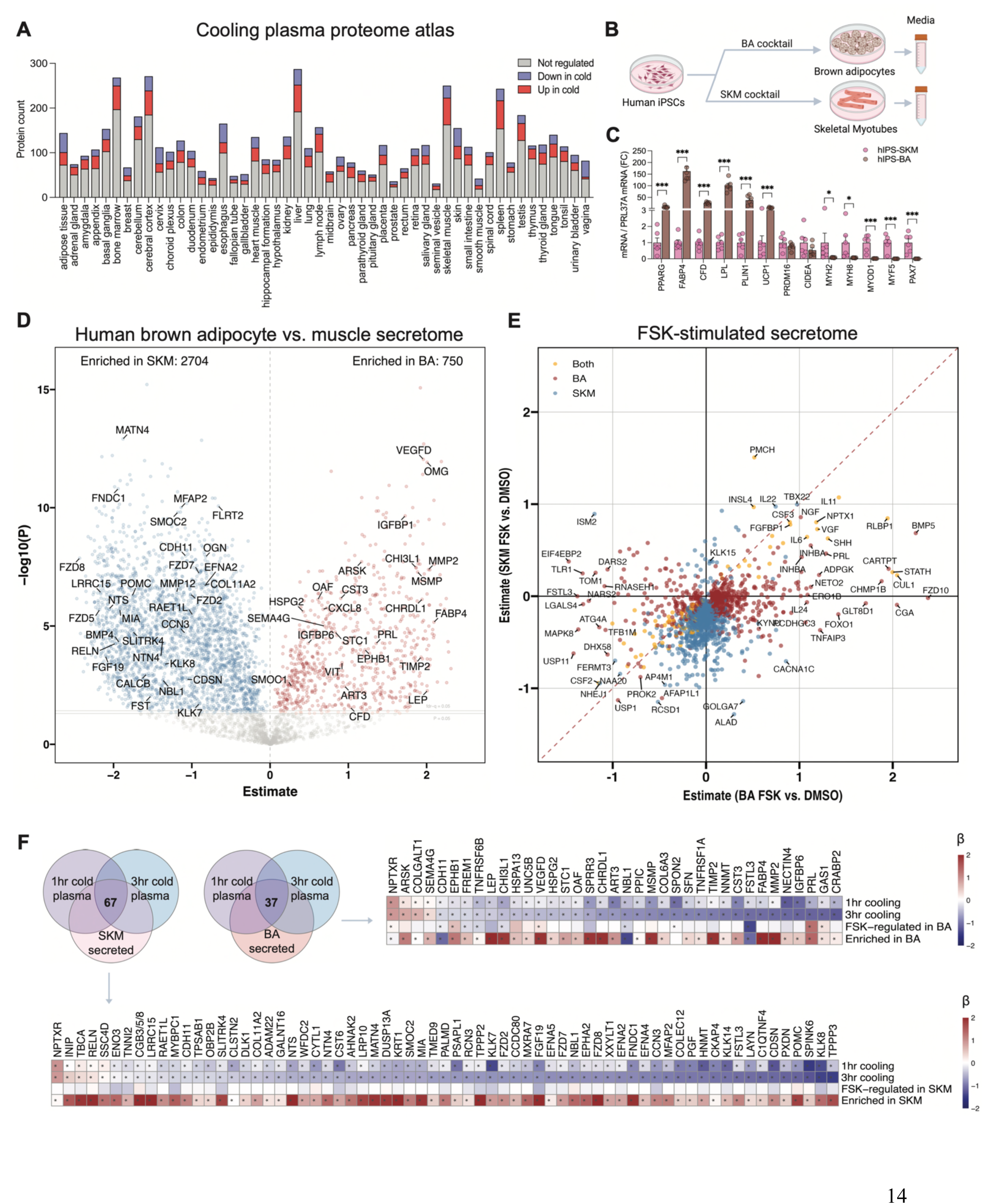
Cold-regulated plasma proteome maps to tissues of origin and brown adipocyte secretome. **A.** Protein source prediction based on Human Protein Atlas. **B.** Experimental setup for identification of secreted factors enriched in human brown adipocytes (hBA) or skeletal muscle cells (hSKM). **C.** Enrichment of adipogenic and thermogenic genes in hBA compared to hSKM cells. Data represent means ± SEM; asterisks: * p < 0.05, *** p < 0.001. See also Figure S2. **D.** Secreted proteomes of mature hBA and hSKM cells. Highlighted are proteins nominally altered with 1hr and 3hr cooling in human plasma. See metadata in Table S2.6 and Figure S2. **E.** Effects of 24hr Forskolin (FSK; 10uM) treatment on the secreted proteomes in hBA and hSKM cells. Plotted are β-estimates for effects of FSK in each cell type. See metadata in Table S2.7. **F.** An overlap between concordantly regulated plasma proteins and factors secreted by human brown adipocytes or skeletal muscle cells. Plotted are proteins that were either enriched in brown adipocyte media (right panel) or skeletal muscle media (bottom panel) or regulated in these cells by forskolin treatment. See also Table S2.8 and S2.9.

### Mapping cold-responsive plasma proteomes to the brown adipocyte secretome

Given that the HPA lacks the brown fat expression profile, and publicly available bulk transcriptomic data from supraclavicular BAT lack cell type-specificity, we turned to *in vitro* models of human brown adipocytes. Here, we utilized human iPSC-derived brown adipocytes (BA) for secretome analysis. Since the same precursors can differentiate into BA^37^ or skeletal muscle cells (SKM)^38^, we differentiated these cells using an adipogenic or myogenic protocol and compared their secreted proteomes on day 41 (Fig. 2B). This allowed us to identify potential human batokines enriched in BA-conditioned media, and SKM-derived myokines enriched in SKM-conditioned media. Following validation of these cells (see differential gene expression in BA vs. SKM, Fig. 2C; UCP1 protein in BA, Fig. S2A; and secreted proteomes PCA, Fig. S2B), we established that several cold-induced plasma proteins were not only secreted by human BA cells, but also significantly enriched compared to SKM media (Fig. 2D; highlighted are cold-regulated plasma proteins; Table S2.6), including ARSK (enrichment β, p=1.92×10^-8^), SEM4G (enrichment β 0.7, p=1×10^-5^), and Oligodendrocyte myelin glycoprotein, OMG (enrichment β 2, p=1.19 × 10^-12^), some of our top concordantly cold-regulated factors. In addition, we observed that while BA cells did not demonstrate a clear enrichment in secreted NPTXR and IL6 at baseline, the concentration of both proteins was significantly induced with forskolin (FSK, mimics cold) (NPTXR: forskolin β 0.12, p<0.001; IL6: forskolin β 1.08, p=1.41×10^-6^; Fig. 2E; Table S2.7), suggesting that brown fat cells secrete these factors upon cold-like stimulation. Overall, the intersection of plasma proteome and cell type-selective secretomes (Fig. 2F; Table S2.8 and S2.9) yielded small but concrete lists of novel potential human batokines such as ARSK, SEMA4G and OMG.

### Cooling suppresses circulating mediators linked with T2D, hypercholesterolemia, hypertension, coronary heart disease and heart failure

We next hypothesized that cooling impacts circulating proteins linked with health and disease, particularly those associated with a cardiometabolic status. To broadly elucidate disease associations, we turned to the newest large-scale plasma proteome atlas of health and disease^39^, comprising a large number of diseases (406 prevalent, 660 incident) and 986 health-related traits in 53,026 individuals (Olink-based, UKBB). Here, we used our 254 concordant cold-regulated proteins, which changed with cooling in the same direction in both major cohorts. To avoid direction bias we included pVal-based proteins in order to obtain a similar number of cold up- and downregulated proteins (see waterfall plot in Fig. 3A). This powerful proteome-phenome resource allowed us to uncover that cooling has a robust impact on circulating proteins linked with major disease outcomes, with a remarkable inverse relationship between the impact of cooling and adverse health conditions (Fig. 3A; Table S2.10). Considering this unbiased list of diseases, we found that cold-suppressed proteins were each linked with up to 190 pathologic conditions, while cold-induced proteins exhibited a much less pronounced association, with multiple proteins in fact being negatively linked with disease ((cut-off: p < 0.05 / (2,920 * 406), where 2,920 is the total number of Olink proteins and 406 is the total number of prevalent diseases)). Careful examination of these disease lists further revealed that cooling suppresses >25 plasma proteins linked with T2D, hypercholesterolemia, hypertension, coronary heart disease, heart failure and liver disease (Fig. 3B; Table S2.11). Using unbiased over-representation analysis (ORA) we further established that indeed the following disease categories were far more robustly linked with the cooling proteome (FDR<0.05): Endocrine and Metabolic, Circulatory, Respiratory and Infectious, both by disease prevalence (Fig. 3C) and incidence (Fig. S3A). Example proteins such as KLK11, APOC1, KLK8 and LY6D and associated disease categories are shown in Fig. 3D-E and Fig. S3B. We did not however find significant associations for these proteins with BAT activity using the available BAT characteristics from our ‘validation cohort’, likely due to low statistical power (n=16 subjects; Fig. S2C).

**Figure 3.**
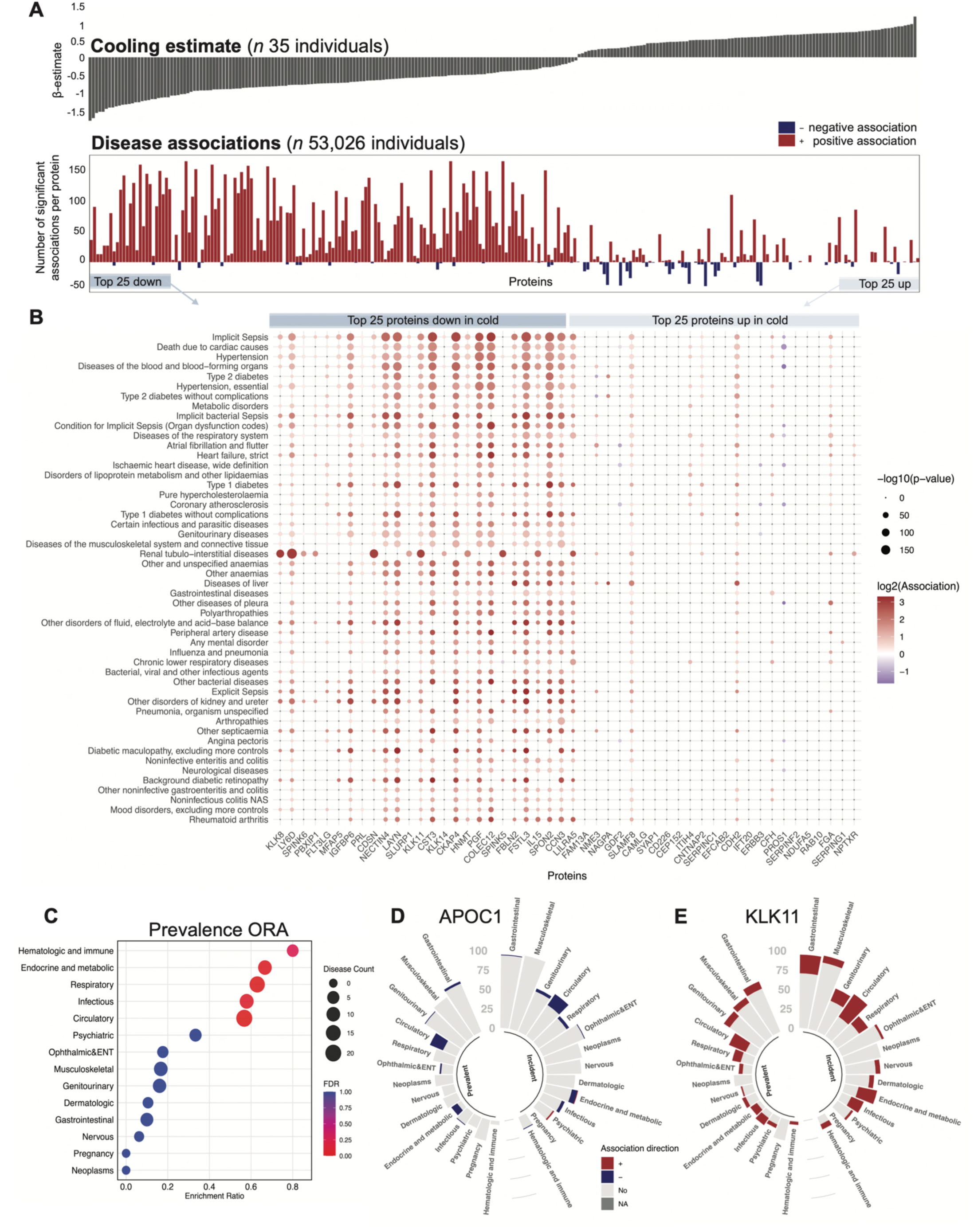
Cold-regulated plasma proteins are associated with health and disease. **A.** Global inverse relationship between the impact of cooling and various disease associations reported in 53,026 individuals (UKBB, Olink). Each bar in the waterfall plot represents a concordant cold-regulated protein, it’s direction of change with cooling (top panel) and number of associated diseases. *Red* indicates positive association between basal protein level and disease, *blue* indicates a negative association between protein level and disease. See metadata in Table S2.10. **B.** Unbiased list of top 50 diseases associated with top 25 proteins decreased or elevated in response to acute cold. See metadata in Table S2.11. **C.** Over-representation of disease chapters in the cold-regulated proteome. Disease data are plotted by prevalence. See incident data in Figure S3. **D.** Example of a cold-induced protein, APOC1 (Apolipoprotein C1), and its negative association with various disease categories by prevalence and incidence. **E.** Example of a cold-repressed protein, KLK11 (Kallikrein 11), and its positive association with various disease categories by prevalence and incidence.

### The circulating signature of cooling resembles a cardioprotective and anti-aging profile

Disease category-specific data, considering top over-represented disease chapters i.e. endocrine and metabolic (Fig. 4A) as well as circulatory diseases (Fig. 4B), further confirmed that T2D, metabolic disorders, hypertension, heart failure, atrial fibrillation and death due to cardiac causes were the most significantly associated health conditions linked with proteins that decrease with cold exposure.

**Figure 4.**
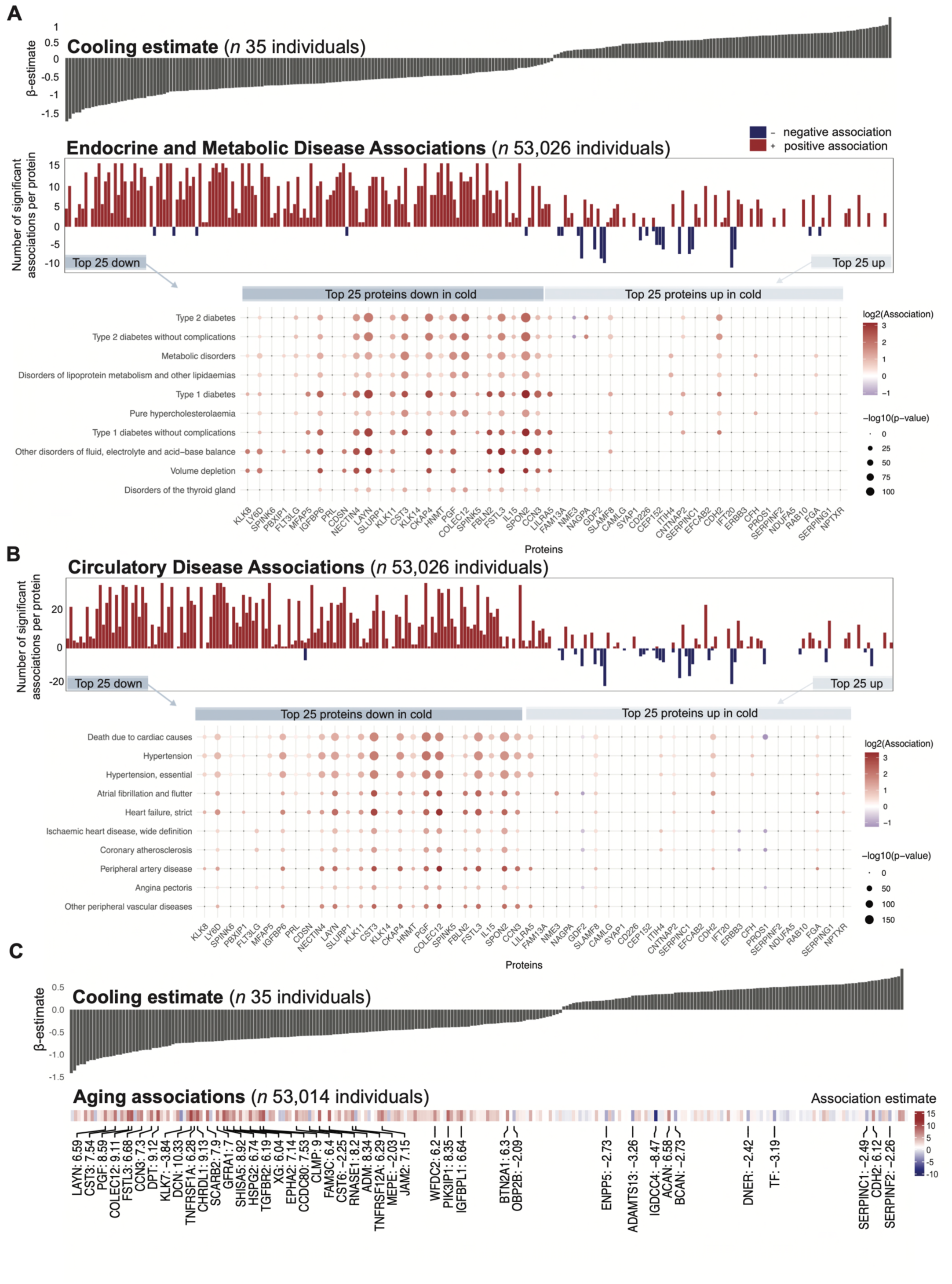
Acute cooling shifts the circulating proteome to a cardioprotective and anti-aging profile in humans. **A.** Phenotypic associations of cold-regulated concordant proteins with Endocrine and Metabolic diseases. Unbiased list of 10 diseases associated with top 25 cold-downregulated and top 25 cold-upregulated proteins. Cooling estimates for corresponding proteins are plotted in the panel above. **B.** Phenotypic associations of cold-regulated concordant proteins (for reference see waterfall plot in panel A) with Circulatory diseases. **C.** Phenotypic associations of 254 cold-regulated concordant proteins with chronological aging in 53,014 individuals (UKBB, Olink). Cooling estimates for corresponding proteins are plotted in the panel above and selected protein names along with mean association strength are indicated in the bottom panel below. See metadata in Table S2.14.

To gain confidence in these findings we examined our concordant list of 254 proteins in an independent UKBB cohort (Plasma Proteomic Profiling; UKBB-PPP), consisting of blood profiles from 54,219 participants^40^. Here we only considered the following phenotypic traits: BMI, fat%, fasting glucose, hypertension (HTN), hyperlipidemia (HLD) and cardiovascular disease (ASCVD). Validating our previous findings, we found that multiple cold-regulated proteins were associated with at least 2 of these traits and exhibited an inverse relationship with cardiometabolic risk factors (Fig. S3C, Table S2.12), i.e. proteins whose levels were suppressed by cooling were overall positively associated with ASCVD, HTN and hyperlipidemia. Among these proteins were cold-suppressed KLK8, LY6D and SPINK6, whose levels were positively associated with BMI, %fat, hyperlipidemia (FDR<0.05), and particularly CVD (p’s<0.0001). In fact, >70% of cold suppressed concordant proteins exhibited positive association with ASCVD, making it the most frequently associated trait in this analysis. In contrast, several proteins whose levels increased with cooling, such as NPTXR exhibited negative associations with ASCVD, HDL and HTN (Fig. S3C). These data suggest that cooling impacts multiple circulating mediators linked with CVD, as well as adiposity and cholesterol metabolism. Interestingly, based on phenotypic associations of predicted pathogenic genetic variants (determined using LoGoFunc^41^; PheWAS) in genes encoding cold-regulated proteins we have additionally found that multiple variants associated with phenotypes based on the 254 concordant list (Table S2.13). For example, a predicted loss-of-function (LoF) variant in COL11A2 is linked with abnormal weight gain. We next focused on cardiometabolic and endocrine phecodes (Fig. S3D-E). Here, we identified several protective effects of risk variants including those linked with total cholesterol, cholesterol/HDL ratio (POMC); cardiac murmurs (LRRK2); and diabetes (RELN).

Finally, since cooling increases metabolic rate and has been proposed to have anti-aging effects due to reduced cellular damage and inflammation^13^, we next examined associations with chronological aging. Here, we utilized longitudinal plasma proteome data based on Olink analysis in 53,014 human subjects in the UKBB to predict age and mortality^42^ and mapped the replicated 254 proteins to the aging association matrix (Table S2.14). Similar to cardiometabolic associations, we saw an inverse relationship between cooling and aging proteomes (Fig. 4C); i.e. multiple cold-suppressed proteins were positively associated with chronological age (e.g. KLK11, SPINK6), while proteins elevated by cooling such as SERPINC1 were linked with a younger age. These data suggest that cooling not only favorably modulates circulating proteins linked with disease but specifically protects against cardiometabolic and age-related pathology.

## DISCUSSION

The study of circulating proteins offers tremendous opportunities to improve diagnosis, risk stratification, and disease prevention. While several endocrine mediators have been proposed to play a crucial role in cold-induced energy metabolism in rodents, plasma proteomic changes in response to cold have largely remained unexplored in humans. Recent technological advances in high-throughput proteomics as well as open access to large-scale proteome-phenome databases have here allowed us to systematically interrogate the circulating proteome linked with acute cold challenge in healthy humans with the goal to uncover cold-sensitive plasma proteins linked with the brown adipocyte secretome, specific cardiometabolic traits, aging and health and disease outcomes.

We measured over 7,000 unique circulating proteins in 94 blood samples and established that about 13% were altered with acute cooling, with 81.5% reproducibility in a validation cohort of BAT+ individuals. Some of the top cold-responsive analytes include known cold-induced proteins such as IL-6 and NPY, but > 95% of the molecules identified here have not been reported in the context of cold exposure in humans.

Unexpectedly, we found that multiple members of Kallikrein-like serine proteases are robustly reduced in the circulation following acute cold adaptation, independent of fasting or CysC/eGFR correction. KLK7 and KLK8 participate in cytokine activation, adipose tissue inflammation, and bradykinin-dependent vasodilation^43, 44^. The robust downregulation of KLK7 and KLK8 during cold exposure may therefore contribute to the anti-inflammatory response, improved insulin sensitivity, and vascular tone modulation during cold exposure. Moreover, adipose-specific deletion of KLK7 was recently demonstrated to prevent mice from gaining WAT mass and sensitized them to insulin, by limiting adipocyte hypertrophy and enhancing browning of subcutaneous depots^44^. Interestingly, based on the UKBB-PPP, we further found that basal KLK8 and KLK11 levels are positively associated with blood glucose, hypertension and CVD in large human populations, while KLK8 was linked with BMI and fat %. The robust downregulation of Kallikreins indeed suggests that this may be a class effect, which likely contributes to improved cardiometabolic health in humans who practice cold exposure.

In fact, over 30% of the cold regulated plasma proteome was linked with both cardiometabolic traits and aging in large human populations. Globally, across cohorts, cooling led to a decrease in proteins that were positively associated with chronological age and cardiometabolic traits such as high BMI, %fat and CVD, such as FABP3, LY6D, SPINK6 and HNMT. Conversely, cooling led to an increase in proteins in the circulation linked to lower odds of these conditions, such as APOE, APOC1 and NPTXR. These data suggest that cooling modulates circulating mediators of metabolic disease, thereby protecting from CVD, T2D, and dyslipidemia. Interestingly, among the top plasma proteins induced with cooling in the two main cohorts, we also found molecules typically annotated as neuronal including SEM4G, NPY, and OMG. Our cell culture data further suggest that a subset of plasma factors identified here are not only secreted by human brown fat cells in culture but also enriched compared to myotubes and potentiated by forskolin treatment, including SEM4G, OMG and ARSK. These data provide a foundation for nominating these proteins as putative BAT-derived endocrine mediators of cold adaptation. Finally, using unbiased proteome-phenome tools, we uncovered for the first time that cooling has a major beneficial impact on proteins associated with adverse health events, particularly those related to endocrine, metabolic and circulatory health, by repressing blood borne proteins linked with diabetes, as well as cardiovascular and liver diseases.

Collectively, our data highlight that the proteomic signature linked with cold exposure, in healthy human subjects, resembles an anti-aging and cardioprotective profile. We provide the scientific community with a reproducible and high-confidence open access resource, with confirmation of BAT specificity using both *in vivo* and *in vitro* methods, to accelerate the development of biomarkers and predictive models for cold therapy-linked prevention of cardiometabolic disease. Our results may also provide previously lacking molecular mechanisms underlying the broad benefits associated with cooling.

## ACKNOWLEDGMENTS

We would like to thank the Rockefeller University Hospital (RUH) team including Rita Devine, Tia Gareau, Lin Zhen, Regina Butler, Jill McCabe, Andrea Ronning, Dacia P. Vasquez, Kadija C. Fofana and Sharon Adams for their help with recruitment and on-site visits. We further extend our thanks to Richard Hutt and Donna Brassil for assistance with Institutional Review Board (IRB) protocols and to Michael Mi, Prashant Rao and Caroline S. Jiang for providing feedback on the analysis. Finally, we would like to thank Jenna Trimboli (RU), and Rosemary Walsh (NIH/NIDDK) for assisting with the three-way institutional agreements.

## AUTHOR CONTRIBUTIONS

K.P. and S.Z. consented study participants and performed cooling/fasting experiments. K.F., C.H., C.P., and L.F. performed the proteomic profiling. L.Y. and K.P. performed cell experiments. K.P. performed gene expression analysis and ELISA validations. Z.C., R.X., T.S.C., G.T and K.P. analyzed data. M.E.K. and Y.I. provided and analyzed PheWAS data. K.Y.C and A.M.C. provided blood samples from two independent human cohorts. R.E.G. provided reagents for Olink and SOMAscan as well as intellectual input throughout the study. O.P., Y.I., A.C. provided intellectual input and supervised parts of analysis. Z.C. and T.S.C. supervised data analysis and statistical comparisons. P.C. and K.P. managed IRB protocols and supervised the research. K.P. and P.C wrote the manuscript with input from all co-authors.

## SUPPLEMENTARY FIGURE LEGENDS

**Figure S1.**
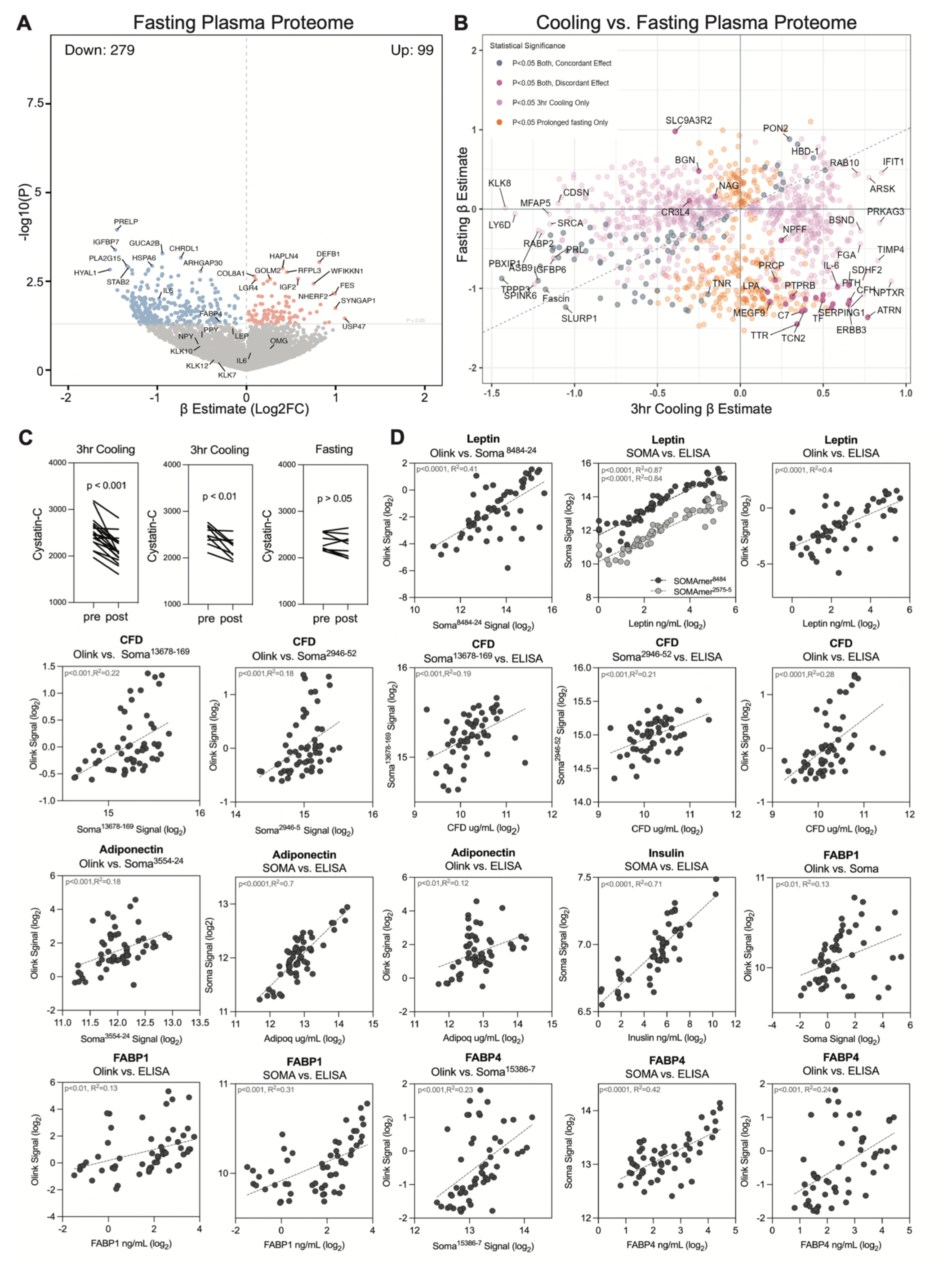
Fasting associated plasma proteome and validations of proteomic data by ELISA. **A.** Effects of fasting on plasma proteins (*n* 9 individuals). See metadata in Table S2.2. **B.** Concordance of cooling and fasting-regulated plasma proteome. **C.** Cystatin C levels before and after cooling or fasting. **D.** Linear regression analysis of individual plasma proteins measured by SOMAScan, Olink and ELISA.

**Figure S2.**
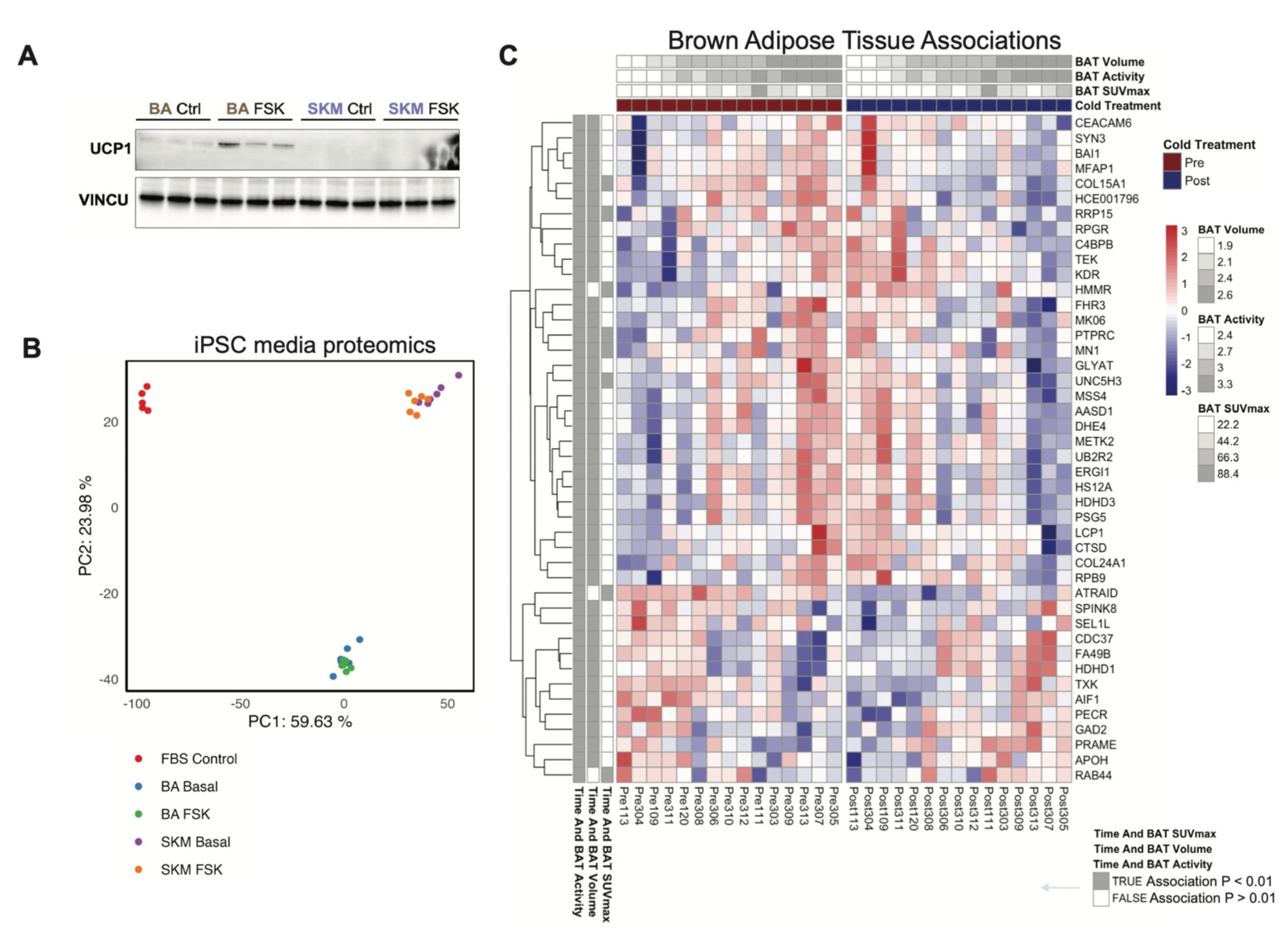
BAT status associations and quality control data for hiPSC-derived cells. **A.** Western blot-based validation of Uncoupling Protein 1 (UCP1) expression in iPS-derived human BA and SKM cell lysates on day 41 of differentiation following Forskolin treatment (10uM, 24hrs). Vinculin is used as a loading control. **B.** Principle component analysis of secreted proteomes in hBA and hSKM cells. **C.** Plasma proteome associations with temperature and BAT traits. Subjects who underwent 1hr cooling are grouped according to their BAT status (volume, activity and SUVmax) from low to high (top panel). Proteins whose expression was altered with cooling dependent on at least 2 of 3 BAT traits (p < 0.01) are plotted (associations with traits are indicated in the left panel; *gray* p < 0.01, *white* p > 0.01).

**Figure S3.**
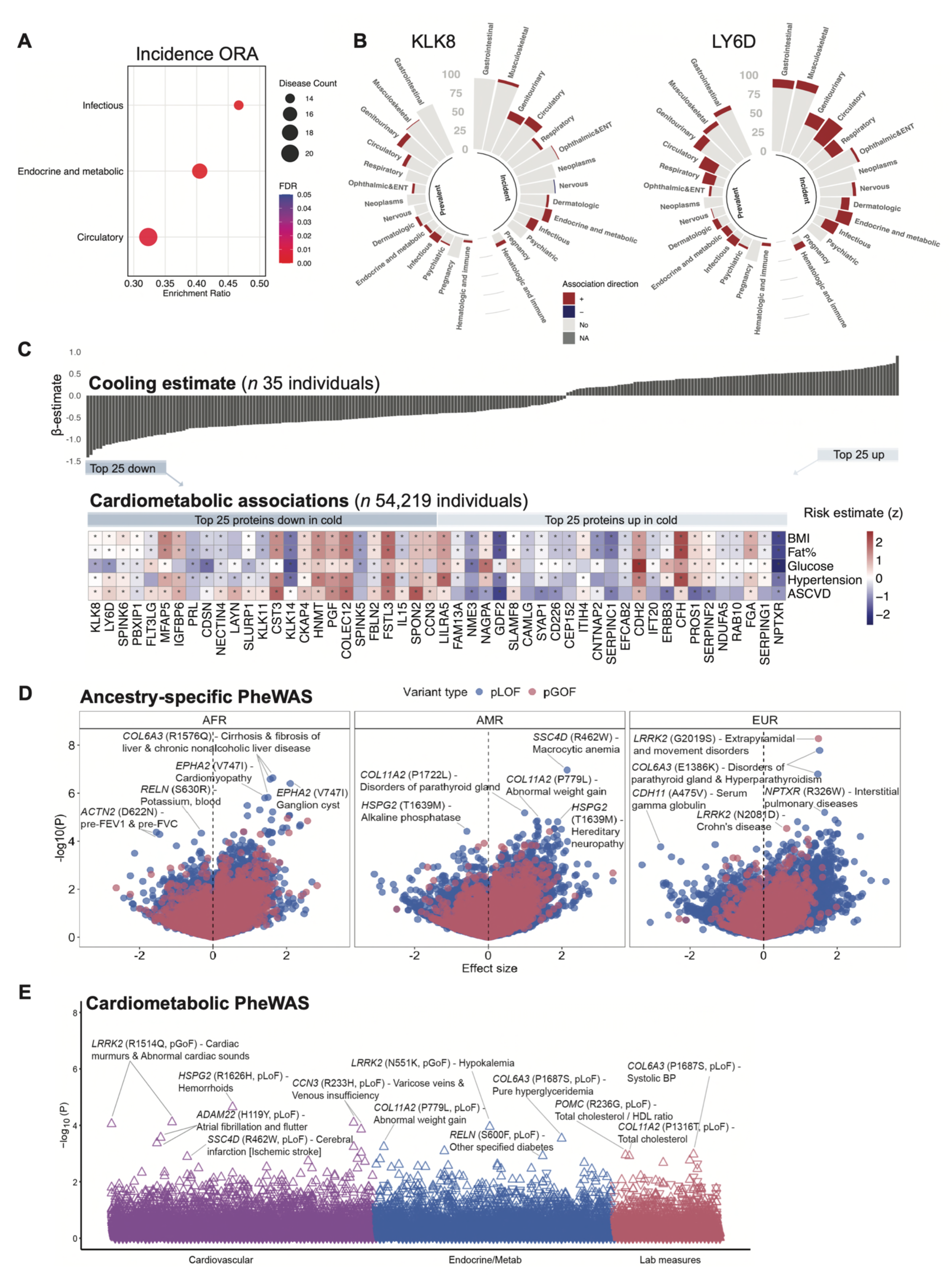
Detailed disease over-representation analysis, cardiometabolic trait associations and PheWAS data. **A.** Over-representation of disease categories in cooling associated plasma proteome plotted by incidence. **B.** Examples of cold-repressed protein KLK8 and LY6D and their positive association with various disease categories by prevalence and incidence. **C.** Phenotypic associations of cold-regulated concordant proteins with cardiometabolic traits based on data from 54,219 individuals with various cardiometabolic phenotypes (UKBB, SOMA). Asterisks indicate associations (FDR<0.05) between basal plasma protein level and a specific trait. Cooling estimates for corresponding proteins are plotted in the panel above and protein names along with mean association strength are indicated in the bottom panel below. See metadata in Table S2.12. **D.** PheWAS of predicted loss (pLoF) or gain of function (pGoF) variants in cold-regulated plasma proteins across individuals of African (AFR, *n* = 13,070), admixed American (AMR, *n* = 9,903), and European (EUR, *n* = 14,745) genetic ancestry in BioMe BioBank. See also Table S2.13. **E.** Manhattan plot showing multi-ancestry meta-analysis in BioMe BioBank (*n* = 37,718) examining cardiometabolic associations of pLOF and pGOF variants in cold regulated plasma proteins. Upward pointing triangles indicate a positive direction of effect, while inverted triangles indicate a negative direction of effect. See also Table S2.13.

**Table S1.** Anthropomorphic characteristics and traits related to cooling procedures in two main cohorts. Data indicate means ± SEM; p values are based on unpaired t-test between groups; *ns* not significant.

**Table S2.** Combined metadata, phenotypic associations and PheWAS of cooling-associated plasma proteome.

## Star ★ METHODS

### KEY RESOURCES TABLE

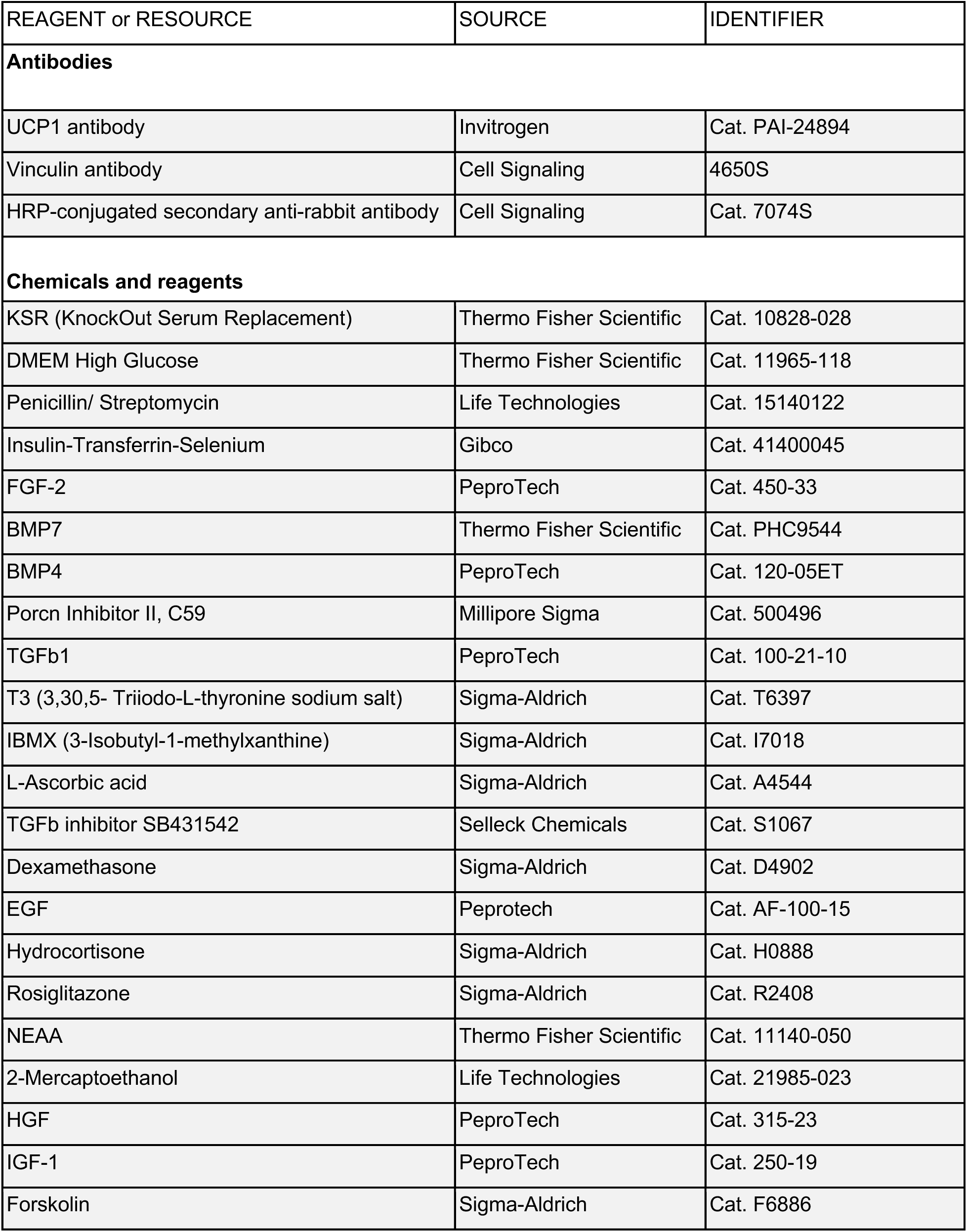
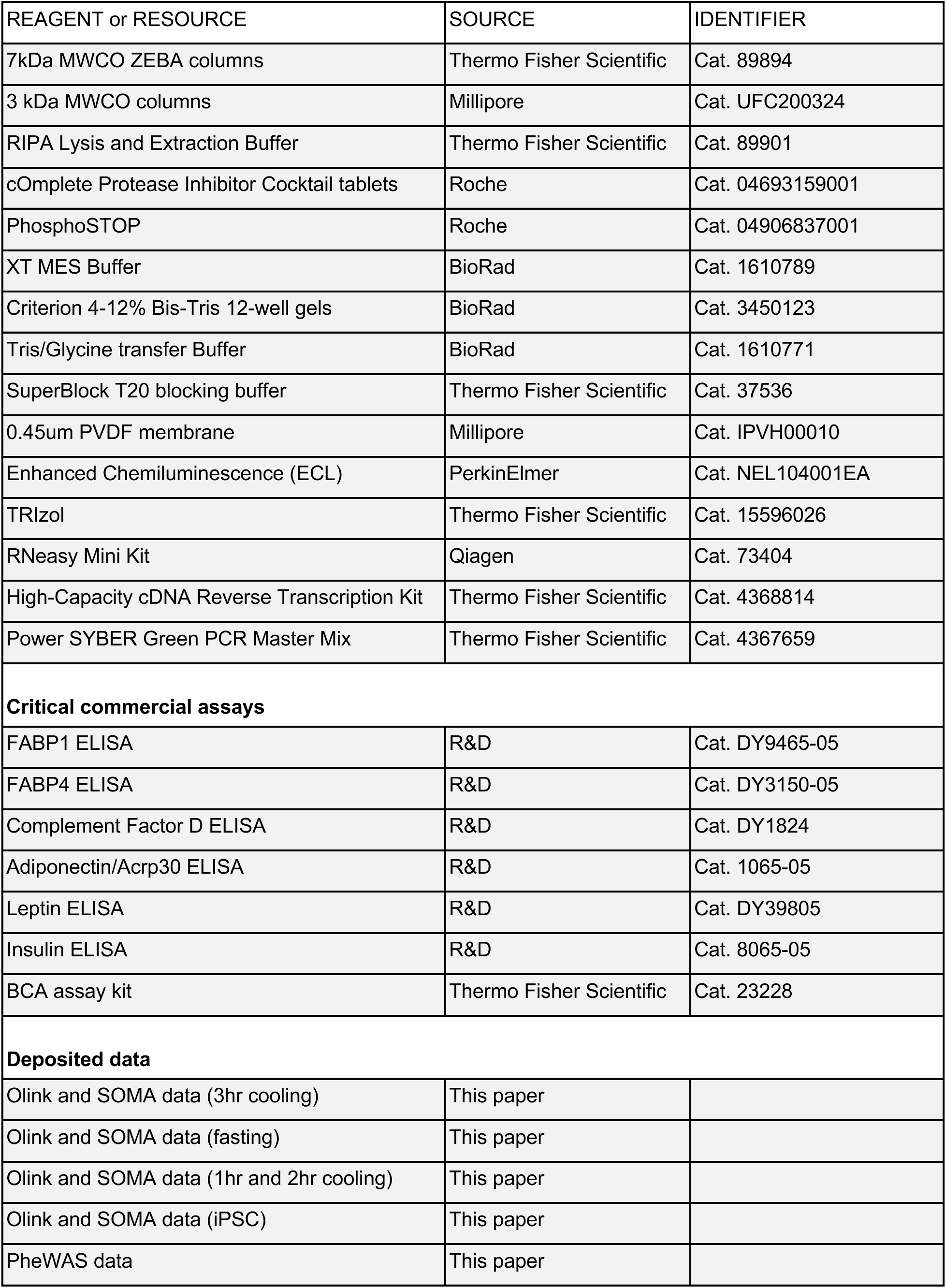
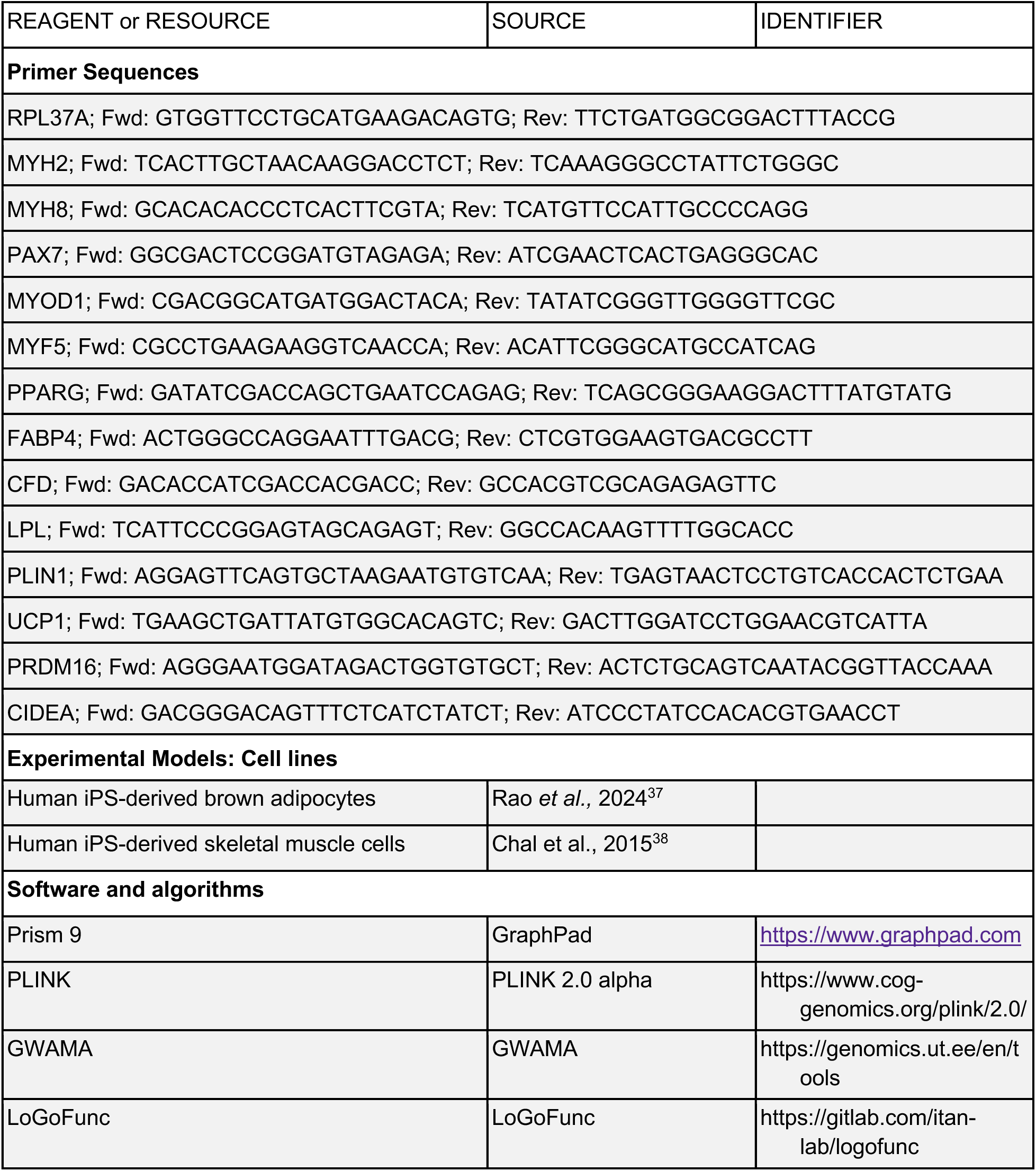

#### Resource Availability

Original code related to LIMMA analysis has been deposited at: https://github.com/RockefellerUniversity/Plucinska_et_al_2025/tree/main

Functional predictions generated with LoGoFunc^42^ are available at https://itanlab.shinyapps.io/goflof/ and https://zenodo.org/records/10126185. Any additional information required to reanalyze the data reported in this paper is available from the lead contact upon request.

#### Subject information and cooling protocol

The study was approved by the Rockefeller University Institutional Review Board (The Cold Vest cohort; clinical trial ID: NCT05050240). All participants provided written informed consent. In total, 20 subjects were recruited for the 3hr cooling experiments at the Rockefeller University Hospital (RUH). Of 20 participants, one was unable to provide blood after cooling and was therefore excluded from the analysis. Proteomics were performed on plasma samples at baseline and after 3 hours of cooling for the remaining 19 participants. All cooling procedures were conducted between October 2021 and February 2022 in New York City when outdoor temperature was on average 8°C and indoor temperature was on average 23°C. Anthropomorphic measurements including height, weight, body mass index (BMI), waist-to-hip ratio (WHR), and body composition measured by air displacement plethysmograph (BodPod; COSMED) on the day of the cooling procedure. Subjects were fasted from 2200hrs until completion of testing the next day (between 1300 and 1400hrs). Blood samples were drawn from the median cubital vein prior to cooling (pre: immediately before cooling, between 0800-0900hrs; at an ambient temperature of 22-24°C; fasted overnight) and immediately after a 3hr personalized cooling protocol (post: 180min after cooling, at an ambient temperature of 18-22°C, fasted overnight + 3 hrs) for paired analysis. For the cooling procedure, participants started at an ambient temperature of 18-22°C. They were then cooled for 3 hours using a water-perfused cooling vest (Polar Products, Stow, OH) attached to a digitally controlled cooler reservoir, which circulates water between the vest and the cooler. The procedure was consistent with BARCIST criteria for human BAT studies^33^. Body temperature was monitored every 15min by tympanic thermometer. During the first 30-45 mins of cooling, individual shivering thresholds (ST) were determined via a ‘shivering’ survey (scores 1-5 were obtained, with 1 = being warm and 5 = being very cold and shivering), while the vest temperature was lowered from 18°C by 1°C every 5 mins. Once an individual’s ST was determined, the vest temperature was adjusted to 2°C above that individual’s ST (corresponding to repeated shivering survey score of 4 – very cold without shivering). All participants were re-warmed after 180min of cooling after the second blood draw was obtained.

All participants were re-invited for a ‘fasting only’ procedure between October and December 2022 (9 to 12 months after the initial cooling procedures), where additional blood draws were collected: fasted 12 hrs and fasted 15 hrs (without the cooling procedure) to control for the effects of fasting. 9 participants (5 males and 4 females) returned for the fasted blood draws (time 0, between 8 and 9am; ambient temperature of 22-24°C fasted overnight) and following 3 hours at RT (time 180 mins, between 1300 and 1400; ambient temperature of 22-24°C) for paired analysis to control for effects of prolonged fasting on plasma proteome.

#### ^18^FDG-PET/CT-confirmed BAT-positive human validation cohorts

Plasma samples from cold-exposed subjects with ^18^FDG-PT/CT scans to detect BAT were obtained from Dr. Aaron Cypess (NIDDK, Bethesda, MD) and are based on two previously published cohorts^31, 32^. From the Baskin study, we obtained plasma samples from 12 men (see Table S1) who underwent 1-hr cold exposure. Blood was drawn at 0800hrs at RT, then all participants entered the cooling chamber with temperature set to 20°C. The cooling vest was turned on at 1200hrs, with temperature set to 15°C and blood was drawn after 1hr cooling (1300hrs, except one individual whose blood draw after 1hr cooling was at 11am). From the O’Mara cohort, we obtained plasma samples from 6 healthy women, who underwent 1-hr and 2-hr cold exposure paired with PET/CT imaging to determine BAT volume and activity (see Table S1). Similar to the male cohort, blood was drawn from the female group at 0800hrs (at RT) prior to cold exposure, then again at 1300hrs and 1400hrs (1 and 2hrs into cooling).

#### Blood collection

Plasma (extracted using K2 EDTA-coated tubes) was obtained immediately following the blood draw and kept frozen in -80°C. Basic clinical blood work was performed at Memorial Sloan Kettering (MSK, New York) and included fasting blood glucose, Hba1c, triglycerides, and TSH (all obtained from time 0 prior to the cooling procedure).

#### Proteomic profiling

Proteomic profiling was conducted on 55ul plasma aliquots using the Soma 7K and 50uL using the Olink 3K platforms run at Beth Israel Deaconess Medical Center (Boston, MA). SomaScan technology leverages the affinity of single-stranded DNA aptamers (‘SOMAmers’) with specific plasma proteins to measure relative protein concentrations in a multiplex reaction. A detailed description of this method is provided in previous publications^45^. In summary circulating target proteins are allowed to bind SOMAmers that are biotinylated and contain a photo-cleavage linker. Bound SOMAmer-protein complexes are captured on streptavidin coated beads, then exposed to conditions that encourage non-cognate SOMAmer-protein complexes to dissociate including a photo-cleavage step. The bound proteins are then biotinylated and recaptured on streptavidin beads, allowing for the isolation of previously bound SOMAmers for quantification using NextGen sequencing. All samples were run in 3 separate batches (batch 1: 3hr cooling Discovery cohort, 2: the fasted controls from the Discovery cohort, and batch 3: the Validation Cohort) with a total of 5,381 common aptamers measured in all cohorts. Intra- and inter-plate variation in measurements were accounted for using normalization and scaling to controls included within each plate. A total of 7,596 aptamers were measured in the cooling and fasting protocols without any missingness. There was no missingness in the SOMA data. The median intra-assay coefficient of variation (CV) was 0.04 in the cooling protocol, 0.04 for the fasting protocol, and 0.03 for the Validation Cohort.

Olink analyses were run on the Olink Explore panel (Olink Proteomics AB) with detailed methods described previously^35, 46^. Briefly, pairs of oligonucleotide-labeled antibody probes bind to a targeted protein. Due to the proximity after binding, oligonucleotide hybridization is then facilitated, allowing for the additional polymerization, extension, and amplification of a unique double-stranded DNA barcode for each specific antigen. The resulting DNA sequence is subsequently detected and quantified using next-generation sequencing (Illumina NovaSeq). Data were quality-controlled and normalized using an internal extension control and a plate control, to adjust for intra- and inter-run variation. The final assay readout was presented in Normalized Protein eXpression (NPX) values (log2-transformed ratio of sample assay counts to extension control counts), where a higher value corresponds to a higher protein expression. Internal controls for incubation, extension, and amplification are included on each plate. Proteomic data were standardized to a set of intraplate control samples and normalized using log transformation. A total of 3,005 proteins were measured using Olink. No proteins had >10% missingness in the cooling protocol, 6 proteins had >10% missing in the fasting protocol in the Discovery Cohort, and 74 proteins in the Validation Cohort. Median intra-assay CVs were 0.34 in the cooling protocol, 0.22 in the fasting protocol, and 0.19 for the Validation Cohort.

#### Validation by ELISA

Six circulating proteins were selected for validation, based on ELISA availability. Paired plasma samples from the cooling procedure and fasted controls were used for determination of FABP1, FABP4, Complement Factor D, Adiponectin/Acrp30, Leptin and Insulin concentrations (see Star Methods Table for details). Plasma dilutions were adjusted to fit the standard range and samples were diluted 1:6 or 1:4 for FABP1, 1:4 for FABP4, 1:10,000 for CFD, 1:15,000 for Adiponectin, 1:20 for Leptin and 1:2 for Insulin. Standards and samples were run in duplicates and optical densities were determined using a microplate reader set to 450 nm with wavelength correction set to 540 nm. Standard curves were created using a four-parameter logistic curve-fit. Protein concentrations were extrapolated and corrected for dilution.

#### Identification of tissue-enhanced plasma proteins

To study the organ specificity of plasma proteins detected in SOMA and Olink platforms, we leveraged the Gene Tissue Expression Atlas (GTEx), a human tissue bulk RNA-seq database. We defined organ-enhanced genes based on at least 4-fold higher mRNA level compared to the average level in all other tissues, in accordance with the definition proposed by the Human Protein Atlas^36^. This resulted in a list of 6,197 tissue-enhanced genes, and their normalized transcript per million (nTPM) across tissues were z-score standardized for the downstream analysis. Out of 7,113 proteins in SOMA and Olink combined dataset of the Cold Vest cohort, 2,366 proteins were mapped to specific tissue(s). For each tissue, the number of tissue-enhanced proteins that respond to cooling was calculated.

#### Human iPS-derived brown adipocytes and skeletal muscle cells

Human pluripotent stem cell-derived brown adipocytes and skeletal muscle cells were differentiated from the same NCRM1 (NIH CRM) human iPS cells as described previously^37, 38^. Briefly, the iPS cells were first differentiated into presomitic mesoderm and then into PAX3/PAX7-expressing dermomyotomal cells which can be propagated to myogenic and non-myogenic lineages. To differentiate into the brown adipocyte lineage, cells were first dissociated and replated on Day 16 of differentiation in DMEM high glucose based medium, supplemented with Penicillin/ Streptomycin, 5% KnockOut Serum Replacement, 1% Insulin-Transferrin-Selenium, 10 ng/ml FGF-2, 10 ng/ml BMP7, 20 ng/ml BMP4 (PeproTech, 120-05ET), 20 nM Porcn Inhibitor II, C59, 10ng/ml TGFb1, 2 nM T3 (3,30,5-Triiodo-L-thyronine sodium salt) for 4 days. Then, the adipocyte precursors were further differentiated in DMEM high glucose, Penicillin/Streptomycin, 10% KnockOut Serum Replacement, 1% Insulin-Transferrin-Selenium, 500 μM IBMX (3-Isobutyl-1- methylxanthine), 25.5 μg/ml L-Ascorbic acid, 2 nM T3, 5 μM TGFb inhibitor SB431542, 1 μM Dexamethasone, 10 ng/ml EGF, 4 μg/ml Hydrocortisone and 1μM Rosiglitazone until day 60. To differentiate into the myogenic lineage, cells were cultured in DMEM high glucose, Penicillin/Streptomycin, 15% KnockOut Serum Replacement, NEAA, 0.01 mM 2-Mercaptoethanol, 10 ng/ml HGF and 2 ng/ml IGF-1. On day 30, the superficial layer was removed mechanically and the progenitors left attached were cultured in the same medium until day 40. Media samples were collected from mature skeletal muscle and brown adipocytes at baseline (24-hr accumulation of secreted molecules) or after a 24-hr Forskolin treatment (FSK, 10uM). 6 mL of media was collected from n=6 cultures per condition. Media samples were de-salted using 7kDa MWCO ZEBA columns, concentrated using 3 kDa MWCO columns and run in SOMA and Olink platforms. DMEM high glucose media supplemented with Penicillin/Streptomycin and KnockOut Serum Replacement were used as negative controls and subjected to profiling in all assays listed above. See Star Methods for reagents details.

#### Gene expression in hiPS cells

RNA from hiPS cell pellets was obtained using a combined TRizol and RNeasy spin column protocol. A total of 1 ug RNA was used for cDNA synthesis per sample. Expression of genes (see Star Methods for primer sequences) was assessed with Sybr Green using QuantStudio 6 Flex Real-Time PCR System in a 384-well plate format. hPRL37a was used a reference gene for delta analysis. Forward and reverse primers sequences are reported in the Star Methods Table. Relative mRNA expression was calculated using the ΔΔCT method with RPL37A as reference gene.

#### Protein expression in hiPS cells

Soluble proteins from hiPS cell pellets were extracted using NP40-based lysis buffer. Lysates were rotated end-over-end for 30 mins at 4°C, spun down at 8,000 rpm, and lipid-free soluble supernatants were extracted. Protein concentrations were determined using a Pierce BCA assay and prepared at 3ug/ul in XT sample buffer, boiled at 90°C for 10 mins and resolved in 12-well 4-12% Criterion XT Bis-Tris gels in MES electrophoresis buffer. Proteins were transferred onto 0.45um ImmobilonP PVDF membranes in Tris-Glycine-MeOH buffer at 30V (4°C, overnight), blocked with SuperBlock for 1hr at RT and probed with anti-UCP1 antibody overnight at 4°C. HRP-conjugated secondary antibody was used for detection with enhanced chemiluminescence in a Chemidoc Imager.

#### Data transformation and paired statistical analyses

Olink NPX values were generated in log2 scale. SOMA proteomics measurements were also log2 normalized. Both protein measurements were z-score transformed within batch. Since both the SOMA and Olink platforms measured circulating proteins, these two datasets were then combined. Of the 7,198 proteins that were measured in total by both platforms, duplicated protein measures were removed and data from one platform for each protein was prioritized based on 1) the presence of a GWAS variant association in the coding region of the protein’s gene and/or 2) having the lower intra-assay covariance (CV). Paired t-tests of proteins measured at time 0hr and 3hr in the Discovery Cohort were used to identify cooling regulated analytes. A Benjamini-Hochberg correction for the total number of analytes measured was used to adjust for multiple hypothesis testing with a false discovery rate (FDR)<0.05 defined as significant^47^. Paired t-tests were also used for proteins measured at baseline after an overnight fast and 3hrs later in the ‘fasting only’ protocol. Paired t-tests were also used to identify changes in analytes with significant differences in response to the cooling protocol (i.e. Time 2 measurement – Time 1 measurement during cooling protocol which is also the log2(FC) since the measurements were in log2 scale that were then z-score standardized within batch) compared to the fasting protocol (i.e. 15hr measurements - 12hr measurements, which were also the log2(FC) and z-score standardized within batch). A similar approach was used to identify circulating proteins that changed with either 1 hour of cold exposure in the Validation Cohort after 1hr cold exposure and 2 hours of cold exposure in a subset of female Validation Cohort individuals.

#### Data residualization

To adjust for the possibility of increased protein consumption, proteolysis, filtration, or hemodilution due to a skew towards decreases in protein levels with cooling, proteins were also residualized to an individual’s Cystatin C levels to adjust for changes in renal filtration or hemodilution. Paired t-tests for CysC as well as CysC-resdiualized data are also presented.

#### LIMMA and correlational analysis with BAT traits

BAT measurements including BAT activity, BAT volume, and maximum SUV (SUVMax) were available for the Baskin and O’Mara studies. Proteomics expression data were imported into R and normalized using variance stabilizing normalization from the VSN package. Following normalization, differential expression analysis was performed using the limma package, incorporating the duplicate correlation function and blocking on subject. The model included Cold treatment, BAT activity, BMI, and SUV.Max as covariates. Proteins showing significant interaction effects between Cold treatment and BAT-related measures were visualized using the heatmap package.

#### Phenome-wide association studies in BioMe BioBank

Phenotypic associations of predicted pathogenic variants in genes encoding cold regulated proteins were investigated in BioMe BioBank. BioMe BioBank comprises exome sequencing (ES) data and electronic health records (EHR) from 37,718 unrelated individuals with African, Admixed American and European genetic ancestries, sequenced at two centers, Regeneron (*n* = 25,369) and Sema4 (*n* = 12,349). Patient diagnoses were extracted from EHR based on ICD-9 and ICD-10 codes, which were then mapped to phecodeX. 1,216 phecodes with at least 50 cases were included in the downstream analyses. Additionally, 381 laboratory measurements and 7 vital signs measured in at least 50 individuals were curated from EHR and normalized using inverse quantile normalization. Predicted pathogenic loss- and gain-of-function (pLoF and pGoF) variants located in genes encoding cold regulated proteins were determined using LoGoFunc^41^. pLoF and pGoF variants with a prediction score > 0.5 and a minor allele count ≥ 20 in each genetic ancestry group were included in the analysis. Single variant association tests were performed in each genetic ancestry group and sequencing cohort (Regeneron and Sema4) separately using Firth’s logistic regression and linear regression for binary and quantitative traits, respectively^48^. Age, sex and first 10 genetic principal components were used as covariates in all analyses. GWAMA was employed to conduct fixed-effects ancestry-specific meta-analyses of Regeneron and Sema4 cohorts^49^. Variants and phenotypes that passed quality control in at least two cohorts were used to perform a random-effects multi-ancestry meta-analysis using GWAMA. *P* values were adjusted according to genomic control (lambda) values in all meta-analyses. and incident diseases, the p values < 10^-5^ were used as the statistical significance threshold.

#### UK BioBank trait and disease association analysis

Aging, disease association and cardiometabolic association UKBB data were obtained from three publicly available datasets^39, 40, 42^. We restricted our analysis to the Olink 3K platform to match their study and generated a list of 254 Olink-specific cooling-associated proteins by overlapping our 1hr and 3hr cooling data (p values < 0.05, same direction of change in response to cold). Disease, cardiometabolic trait and aging associations for cooling-regulated proteins were extracted and ordered by 3hr cooling estimates for visualization. We used very stringent cut-offs for disease association data, based on the number of comparisons, i.e. Prevalence (406 prevalent diseases), hence p < 0.05 / (2,920 * 406), where 2,920 is the total number of proteins detected by Olink; Incidence (660 incident-based diseases), hence p < 0.05 / (2,920 * 660). To identify disease categories linked with cooling-regulated proteome, we performed over-representation analysis on top 100 diseases that have the greatest number of associations per protein. We used all the diseases in International Classification of Diseases (ICD)-10 codes as background list, which classifies diseases into 14 different chapters. For each disease chapter, a one-tailed hypergeometric test was used to calculate the probability of observing greater number of diseases from that chapter in the selected set, given its frequency in the background. P values were adjusted for multiple comparisons using the Bonferroni method.

