## Supplementary Table 1 for "Acute cold exposure in humans shifts the circulating proteome to a cardioprotective and anti-aging profile"

**Table S1.** Anthropomorphic characteristics, traits related to cooling procedures and brown adipose tissue in Discovery and Validation cohorts. Data indicate means ± SEM; p values are based on unpaired t-tests between groups; *ns* not significant.

| **Subject characteristics** | **Discovery cohort** | | | |
|---|---|---|---|---|
| | Males | Females | t-test | Combined |
| n | 9 | 10 | - | 19 |
| Age | 24.4 ± 0.7 | 25.1± 0.6 | ns | 24.8 ± 0.4 |
| Height | 176.8 ± 2.2 | 164.1 ± 2.2 | p < 0.001 | 170.1 ± 2.1 |
| Mass (kg) | 67.7 ± 2.5 | 59.5 ± 2.3 | p < 0.05 | 63.4 ± 1.9 |
| BMI (kg/m$^2$) | 21.6 ± 0.6 | 22.1 ± 0.5 | ns | 21.9 ± 0.4 |
| WHR | 0.85 | 0.75 | p < 0.001 | 0.8 |
| Fat (%) | 14.5 ± 2.9 | 28.1 ± 2 | p < 0.001 | 21.6 ± 2.3 |
| Fat-free mass (%) | 85.5 ± 2.9 | 72.0 ± 2 | p < 0.001 | 78.4 ± 2.3 |
| Body Density (kg/L) | 1.06 ± 0.006 | 1.04 ± 0.004 | p < 0.001 | 1.05 |
| Fasting glucose (mg/dl) | 92.3 ± 3.2 | 89.7 ± 1.5 | ns | 90.9 ± 1.7 |
| Hba1c (%) | 5.14 ± 0.1 | 4.98 ± 0.1 | ns | 5.06 ± 0.01 |
| TG (mg/dl) | 52.5 ± 4.7 | 84.3 ± 18.4 | ns | 70.2 ±10.9 |
| TSH (mIU/L) | 1.9 ± 0.2 | 1.4 ± 0.2 | ns | 1.6 ± 0.1 |
| Latency to Shiver (min) | 57.7 ± 5.7 | 50.9 ± 11 | ns | 54.3 ± 6.1 |
| Shivering Threshold (ºC) | 9.2 ± 0.6 | 11.0 ± 0.9 | ns | 10.2 ± 0.6 |
| | **Validation cohort** | | | |
| | Males | Females | t-test | Combined |
| n | 12 | 6 | - | 18 |
| Age | 24.4 ± 1.2 | 28.0 ± 1.8 | ns | 25.9 ± 1.1 |
| Height | 183.2 ± 1.5 | 163.0 ± 2.1 | p < 0.001 | 175.1 ± 2.6 |
| Mass (kg) | 78.9 ± 2.6 | 70.6 ± 4.6 | ns | 75.6 ± 2.5 |
| BMI (kg/m$^2$) | 23.5 ± 0.5 | 26.8 ± 1.8 | p < 0.05 | 24.8 ± 0.8 |
| WHR | 0.81 ± 0.01 | 0.82 ± 0.02 | ns | 0.81 ± 0.01 |
| Fat (%) | 16.4 ± 1.5 | 36.9 ± 2.3 | p < 0.001 | 24.6 ± 2.6 |
| Lean mass (%) | 79.4 ± 1.4 | 60.9 ± 2.2 | p < 0.001 | 71.9 ± 2.4 |
| Axial BAT activity | 1319.3 ± 265.7 | 1141.9 ± 491.8 | ns | 1260.2 ± 233.6 |
| Axial BAT volume (mL) | 307.9 ± 44 | 183.7 ± 49.7 | ns | 266.5 ± 35.8 |
| Axial BAT SUV Max | 35.4 ± 4.6 | 35.5 ± 14.5 | ns | 35.5 ± 5.5 |
| Fasting glucose (mg/dl) | 90.1 ± 1.4 | 89.5 ± 1.5 | ns | 89.8 ± 0.9 |
| Insulin | 7.8 ± 1.3 | 10.6 ± 1.5 | ns | 8.9 ± 1 |
| HOMA-IR | 31.6 ± 5.6 | 42.1 ± 6.3 | ns | 35.9 ± 4.3 |
